# Activation of hTREK-1 by polyunsaturated fatty acids does not only involve membrane tension

**DOI:** 10.1101/2022.08.01.502268

**Authors:** Emilie Bechard, Elodie Arel, Jamie Bride, Julien Louradour, Xavier Bussy, Anis Elloumi, Claire Vigor, Pierre Soule, Camille Oger, Jean-Marie Galano, Thierry Durand, Jean-Yves Le Guennec, Hamid Moha-Ou-Maati, Marie Demion

**Affiliations:** PhyMedExp, Université de Montpellier, Inserm U1046, UMR CNRS 9412, Montpellier (France); IBMM, Université de Montpellier, UMR CNRS 5247, ENSCM, Montpellier (France); IGF, Université de Montpellier, Inserm 1191, UMR CNRS 5203, Montpellier (France) Present adress INM, Inserm U1298, Montpellier (France); NanoTemper Technologies GmbH, Munich (Germany)

**Author notes:** last co-authors (Jean-Marie Galano, Thierry Durand, Jean-Yves Le Guennec, Hamid Moha-Ou-Maati, Marie Demion). corresponding author Marie DEMION PhyMedExp, Inserm U1046, CNRS UMR9214 CHU Arnaud de Villeneuve Bâtiment Craste de Paulet 370 avenue du Doyen Gaston Giraud 34290 MONTPELLIER Cedex 05 France.

## Abstract

TREK-1 is a mechanosensitive channel also activated by polyunsaturated fatty acids (PUFAs). In this study, we compared the effect of multiple fatty acids and ML402. First, we showed a variable TREK-1 activation by PUFAs related to the variable constitutive activity of TREK-1. Then, we observed no correlation between TREK-1 activation and acyl chain length or number of double bonds suggesting that the bilayer-couple hypothesis cannot explain by itself the activation of TREK-1 by PUFAs. The membrane fluidity measurement is not modified by PUFAs at 10 µM. The spectral shift analysis in TREK-1-enriched microsomes indicates a K_D,TREK1_ at 44 µM of C22:6 n-3. PUFAs display the same activation and reversible kinetics than the direct activator ML402 and activate TREK-1 in both whole-cell and inside-out configurations of patch-clamp suggesting that the binding site of PUFAs is accessible from both sides of the membrane, as for ML402. Finally, we proposed a two steps mechanism for TREK-1 activation by PUFAs: first, insertion into the membrane, without fluidity or curvature modifications, and then interaction with TREK-1 channel to open it.

## INTRODUCTION

Potassium channels have a crucial role in the electrical activity of excitable cells such as neurons and cardiomyocytes. Some of these potassium channels are common drug targets modulating the action potential shape and consecutively organs functions. Among the K^+^ channels, members of the two-pore domains K^+^ channels family (K2P) are involved in the repolarization phase of action potential and in the resting membrane potential (Kelly *et al*, 2006). K2P family includes 15 members classified in 6 functional subfamilies: TWIK (Tandem of pore domains in a Weak Inward rectifying K channel), TREK (TWIK-Related K channel), TASK (TWIK-related Acid Sensitive K channel), TALK (TWIK-related Alkaline pH-activated K channel), THIK (Tandem pore domain Halothane-Inhibited K channel), TRESK (TWIK-Related Spinal Cord K Channel). K2P channels share common structural features, each subunit containing two-pore domains (P1, P2) and four putative transmembrane segments (M1-M4) (Honoré, 2007). The dimerization of K2P channels allows the formation of the canonical K^+^ selective pore domain. TREK-1 channel, one of the three members of the TREK subfamily with TREK-2 and TRAAK, has been discovered in 1996 by Fink *et al*. The gating of TREK-1 channel is poly-modulated by a wide range of physical and chemical stimuli including mechanical stretch, temperature, voltage, pH changes, pharmacological agents and polyunsaturated fatty acids (PUFAs). This channel is widely studied since its activation is involved in neuroprotection (Lamas & Fernández-Fernández, 2019), cardioprotection (Kamatham *et al*, 2019), analgesia (Li & Toyoda, 2015) and reduced epilepsy crisis (Heurteaux *et al*, 2004). In the majority of these diseases, the protection afforded by TREK-1 activation is due to the hyperpolarization of the membrane potential (Djillani *et al*, 2019)

Among the wide diversity of TREK-1 modulators, PUFAs have been shown to behave as strong activators (Patel *et al*, 1998; Danthi *et al*, 2003). PUFAs are amphipathic molecules with a hydrophilic carboxyl head and a long hydrophobic chain of carbons and multiple double bonds. The two main classes of PUFAs are n-3-PUFAs and n-6-PUFAs based on the position of the first double bond from the carbon ω starting at the methyl extremity. Numerous studies suggest that n-3-PUFAs, such as docosahexaenoic acid (DHA) and eicosapentaenoic acid (EPA), exert antiarrhythmic properties through the modulation of ionic channels and consequent membrane hyperpolarization (Kang & Leaf, 1994, 1996). Such hyperpolarization could be consecutive to TREK-1 activation as it is largely expressed in the myocarde(Wiedmann *et al*, 2021; Bechard *et al*, 2022). Since TREK channels are mechano-sensitives, the comparison of the effects of different PUFAs on TRAAK (Fink *et al*, 1996) suggests that PUFAs can insert inside the membrane inducing an increase in membrane fluidity (Leifert *et al*, 1999) that in turn modifies membrane curvature and tension (Sheetz and Singer., 1974). In fact, TRAAK is very likely to be activated by PUFAs through this mechanosensitive pathway since the activation is related to the acyl chain length and to the number of double bonds (Fink et al., 1998; Maingret *et al*, 1999; Patel et al., 2001; Honoré, 2007). However, while it is known that PUFAs also activate TREK-1 channel, no study has compared the effects of different PUFAs on TREK-1 to determine a potential mechanism of action. Here, we propose a study that allows a better understanding of which features of PUFAs are essential for their effects on TREK-1. We performed a thorough comparison of the effects of 9 PUFAs (from 18 to 22 carbons with 2 to 6 double bonds) and 2 other C18 FA (mono-unsaturated and saturated) on the TREK-1 current (I_TREK-1_). We report that there is no correlation between the acyl chain length and TREK-1 activation, as for the PUFAs-induced membrane fluidity. Then, the comparison of the effects of PUFAs and ML402, a direct activator of TREK-1 binding within a cryptic pocket behind the selectivity filter (Lolicato *et al*, 2017), suggests that there is a at least one binding site for PUFAs on TREK-1 channel. This hypothesis is reinforced by the affinity protein-PUFA test performed in TREK-1 enriched-microsomes using the nanotemper technology, but the precise binding site remains to be determined in further studies.

## RESULTS

### TREK-1 channel activation by PUFAs does not depend of acyl chain length

As previously described,TREK-1 channel is activated by PUFAs, such as arachidonic acid (AA, C20:4 n-6) (Patel *et al*, 1998), LA (Danthi *et al*, 2003) or DHA (Ma & Lewis, 2020). In our hands, the average membrane potential (Em) of HEK hTREK-1 cells was −69.2 ± 0.7 mV (mean ± SEM) (**Fig 1D**, n=130), with a variable initial current density of 15.3 ± 1.2 pA/pF (n=130) at 0 mV. During PUFA superfusion, the current density elicited by voltage ramp from −100 mV to +30 mV progressively increased until a steady-state was reached (**Fig 1A and 1B**). When TREK-1 is activated, Em hyperpolarized to −81.7 ± 0.3 mV (**Fig 1D**, n=130), close to the theorical equilibrium potential of K^+^ ions (E_K_ = −86.5 mV). In order to determine wether the variability of the initial current density (I_0_) was related to a variability of TREK-1 channel activity in initial condition, we superfused an inhibitor of TREK-1 channel, Norfluoxetine (NrFlx) at 10 µM. As the initial current was significantly decrease from 8.8 ± 2.2 pA/pF to 3.7 ± 0.9 pA/pF (p-value=0.02; n=5) when Norfluoxetine was applied and as the characteristic outward rectification of I_TREK-1_ was lost, we concluded that I_0_ was carried out mostly by TREK-1 channel (**Fig 1C**).

**Figure 1:**
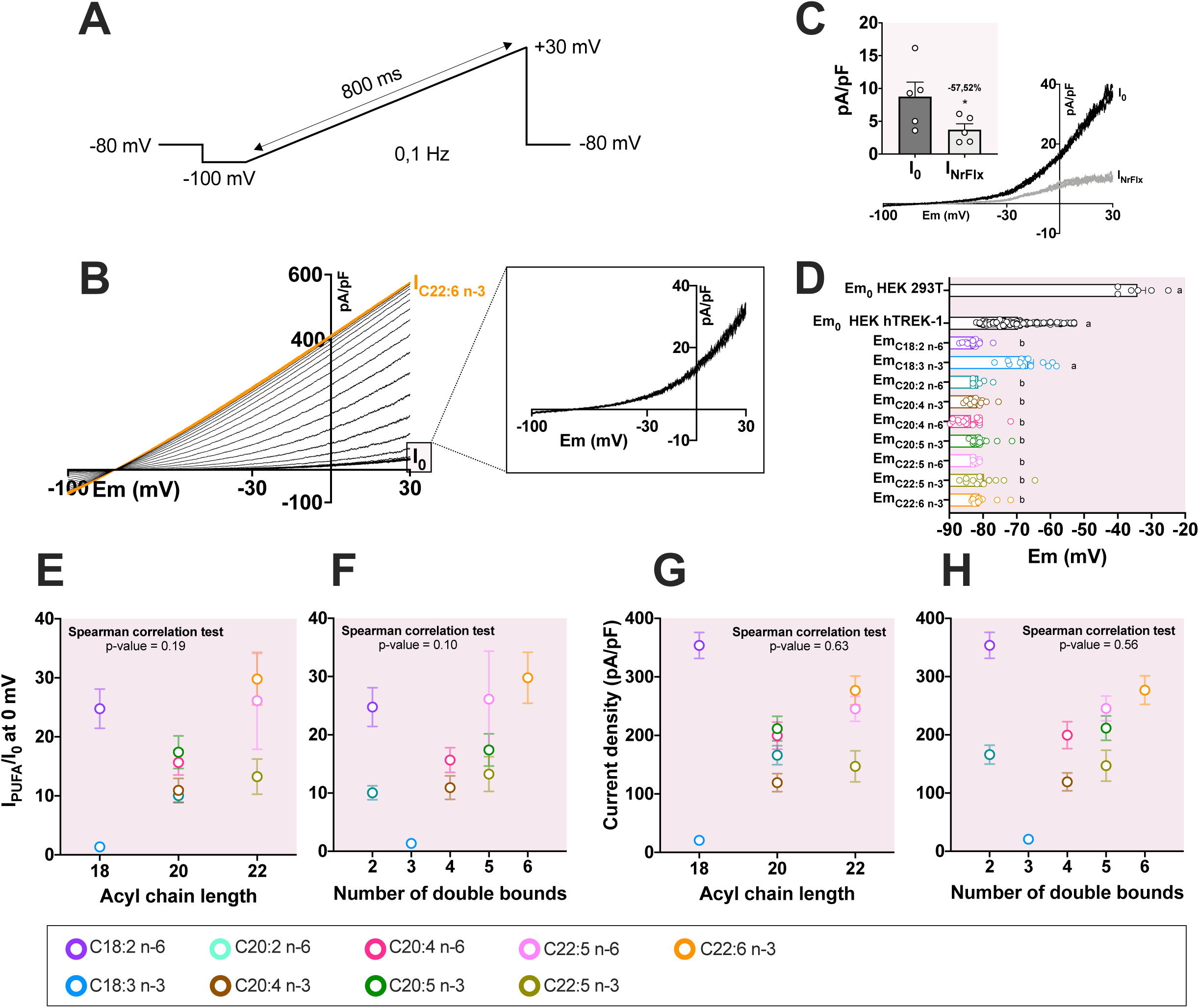
Activation of TREK-1 channel by PolyUnsaturated Fatty Acids. A. Illustration of the voltage ramp protocol to record I_TREK-1_ between −100 mV and +30 mV. **B.** A representative traces of current recordings during C22:6 n-3 (DHA) application is shown in one trace *per* 10s (middle). The inset on the right shows the characteristic outward rectifying initial current of TREK-1 (I_0_: initial current density). **C.** Effect of 10µM of Norfluoxetine (NrFlx on I_0_. **D.** Bar graph showing the initial membrane potential Em_0_ (mV) for HEK 293T and HEK hTREK-1 cell lines and the membrane potential Em_PUFA_ of HEK hTREK-1 cells after the PUFA perfusion. **E.** Lack of correlation between the acyl chain length and I/I_0_ and **F.** the current density (pA/pF), in response to 10 µM PUFAs at 0 mV. **G.** Lack of correlation between the number of double bounds and I/I_0_ and **H.** the current density (pA/pF), in response to 10 µM PUFAs at 0 mV. (Purple: C18:2 n-6, blue: C18:3 n-3, light green: C20:2 n-6, brown: C20:4 n-3, red: C20:4 n-6, green: C20:5 n-3, light green: C22:5 n-3, pink: C22:5 n-6, orange: C22:6 n-3).

To investigate the importance of the acyl chain length and double bounds, we compared the response of 9 PUFAs with various chain lengths, from 18 to 22 carbons with different number of double bounds, from 2 to 6 unsaturations (**Table 1**). We plotted the relationship between current density at 0 mV in the presence of PUFA (I_PUFA_) normalized to the initial current density (I_0_)(this normalized parameter corresponds to the current fold-increase (I_PUFA_/I_0_)) and the number of carbons on the acyl chain. As shown in **Fig 1E**, there is no correlation between PUFA activation I_PUFA_/I_0_ and the acyl chain length (Spearman correlation test: p-value = 0.19), as for the current density of TREK-1 channel after PUFA perfusion (Spearman correlation test: p-value = 0.10) (**Fig 1G**). We then plotted the relationship between I_PUFA_/I_0_ at 0 mV or current density in response of PUFA perfusion and the number of double bounds in the acyl chains and also observed no correlation (**Fig 1F and Fig 1H**; Spearman correlation tests: p-value = 0.63 and p-value = 0.56 respectively). These data reveal that there is no relationship between the effect of PUFAs on TREK-1 and the acyl chain length or the number of double bonds. Indeed, C22:6 n-3 tends to have a stronger effect than C22:5 n-3 on TREK-1 current, while they share the same number of carbons (**Fig 1E-H** and **Table 1**). Accordingly, the most potent activators were both one of the shortest one, C18:2 n-6 (I_PUFA_/I_0_ = 24.8 ± 3.3; current density = 353.7 ± 22.4 pA/pF), and one of the longest one, C22:6 n-3 (I_PUFA_/I_0_ = 29.8 ± 4.4; current density = 276.8 ± 24.5 pA/pF). Conversely, C18:2 n-6 is one of the most potent activator but C18:3 n-3 failed to activate TREK-1 channel (I_PUFA_/I_0_ = 1.4 ± 0.1; Current density = 20.7 ± 4.2 pA/pF). Like C18:3 n-3, the saturated stearic acid (C18:0) had no effect on TREK-1 while the mono-unsaturated C18:1 n-9 produced a 7.4 ± 1.9-fold increase of I_TREK-1_ (n=12, **Table 1**). However, statistical analysis failed to discriminate the PUFA’s effects (I_PUFA_/I_0_ parameters compared with a nonparametric kruskall-wallis test) probably due to the important variability of the effects.

**Table 1:** Fatty acids nomenclature, characteristics and effects on the TREK-1 current in whole-cell configuration of patch-clamp. Values of I_PUFA_/I_0_ and I_PUFA_ are expressed as mean ± SEM.

| | IUPAC Name | | Tail length | Unsaturation number | $\omega$ | $I_{PUFA}/I_0$ | $I_{PUFA}$ (pA/pF) |
| --- | --- | --- | --- | --- | --- | --- | --- |
| **C18:0** | Stearic acid | 18:0 | 18 | 0 | | $1.0 \pm 0.1$ | $18.4 \pm 5.5$ |
| **C18:1 n-9** | Oleic acid (9) | 18:1 $\Delta$ 9 | 18 | 1 | ω-9 | $7.4 \pm 1.9$ | $113.1 \pm 24.6$ |
| **C18:2 n-6** | Linoleic acid (9,12) | 18:2 $\Delta$ 9,12 | 18 | 2 | ω-6 | $24.8 \pm 3.3$ | $353.7 \pm 22.4$ |
| **C18:3 n-3** | α-Linolenic acid (9,12,15) | 18:3 $\Delta$ 9,12,15 | 18 | 3 | ω-3 | $1.4 \pm 0.1$ | $20.72 \pm 4.2$ |
| **C20:2 n-6** | Eicosadienoic acid (11,14) | 20:2 $\Delta$ 11,14 | 20 | 2 | ω-6 | $10.1 \pm 1.2$ | $166.1 \pm 16.2$ |
| **C20:4 n-3** | Eicosatetraenoic acid (8,11,14,17) | 20:4 $\Delta$ 8,11,14,17 | 20 | 4 | ω-3 | $10.9 \pm 2.0$ | $119.2 \pm 15.5$ |
| **C20:4 n-6** | Eicosatetraenoic acid (5,8,11,14) | 20:4 $\Delta$ 5,8,11,14 | 20 | 4 | ω-6 | $15.7 \pm 2.1$ | $199.5 \pm 23.1$ |
| **C20:5 n-3** | Eicosapentaenoic acid (5,8,11,14,17) | 20:5 $\Delta$ 5,8,11,14,17 | 20 | 5 | ω-3 | $17.4 \pm 2.8$ | $211.7 \pm 21.0$ |
| **C22:5 n-6** | Docosapentaenoic acid (4,7,10,13,16) | 22:5 $\Delta$ 4,7,10,13,16 | 22 | 5 | ω-6 | $26.1 \pm 8.3$ | $245.3 \pm 21.4$ |
| **C22:5 n-3** | Docosapentaenoic acid (7,10,13,16,19) | 22:5 $\Delta$ 7,10,13,16,19 | 22 | 5 | ω-3 | $13.3 \pm 2.9$ | $147.1 \pm 26.6$ |
| **C22:6 n-3** | Docosahexaenoic acid (4,7,10,13,16,19) | 22:6 $\Delta$ 4,7,10,13,16,19 | 22 | 6 | ω-3 | $29.8 \pm 4.4$ | $276.8 \pm 24.5$ |

**The variability in PUFA responses is related to the variable initial current density** Despite an important number of cells studied, we observed a large variability of the TREK-1 current activation by PUFAs illustrated in **Fig 2A**. The severity of the inclusion criteria (see Material and Methods section) suggests that the variability observed in PUFAs responses is inherent to TREK-1 channel.

**Figure 2:**
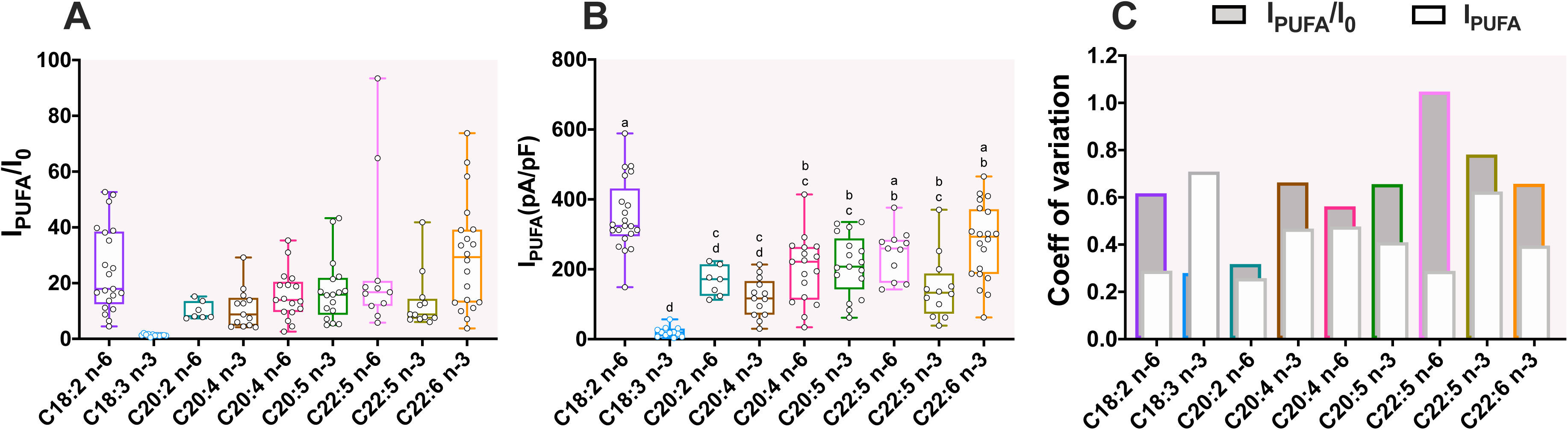
Variability of TREK-1 activation by PUFAs. A. Boxplot of the fold increase (I_PUFA_/I_0_) of TREK-1 and **B**. the current density at steady-state (I_PUFA_, pA/pF) at 0 mV for each PUFA at 10 µM. Boxplots represent the median (lines) with max and min values (error bars). Groups were compared with a Kruskal-Wallis test followed with the *post-hoc* Dunn’s test. Two bars having the same letter are not significantly different. **C.** Changes in the coefficient of variation (SD/mean) between I_PUFA_/I_0_ and I_PUFA_ at 0 (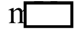: coefficient of variation for I_PUFA_/I_0_;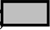: coefficient of variation for I_PUFA_). Both coefficient of variation are superimposed.

**Fig 2A** illustrates the large variability of PUFA effects based on the fold-increase analysis (I_PUFA_/I_0_).We demonstrated that the variability of the I_PUFA_/I_0_ parameter resulted from the variability of the initial current I_0_ but not from the current density at steady-state after the application of the PUFAs (I_PUFA_) (**Fig 2B**). Indeed, the calculation of the coefficient of variation (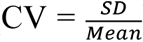) confirmed that the dispersion of the I_PUFA_/I_0_ calculation is higher than the dispersion of the I_PUFA_ at steady-state of the activation (**Fig 2C**). Thus, CV modification is in accordance with the hypothesis that the variability of I_0_ is responsible of the variability of the I_PUFA_/I_0_ parameter.

To better characterize the relationship between I_0_ and I_PUFA_/I_0_, we plotted the fold-increase of TREK-1 current (I_PUFA_/I_0_, Y axis) as a function of the initial current (I_0_, X axis). As shown in **Fig 3A**, there is a non-linear relationship between I_PUFA_/I_0_ and I_0_. This relationship can be linearized by log-transforming the data (Log10) (**Fig 3B**). Thus, the effects of all PUFAs but C22:5 n-3 depends on I_0_, independently of the absolute amplitude of I_PUFA_. Therefore, there is a negative relationship between Log10(I_PUFA_/I_0_) and Log10(I_0_): C18:2 n-6 (R^2^=0.82, p-value=0.0001), C20:2 n-6 (R^2^=0.66, p-value=0.0271), C20:4 n-3 (R^2^=0.80, p-value=0.0001), C20:4 n-6 (R^2^=0.50, p-value=0.0023), C20:5 n-3 (R^2^=0.53, 0.0008), C22:5 n-6 (R^2^=0.84, p-value=0.0001) and C22:6 n-3 (R^2^=0.68, p-value=0.0001) (**Fig 3B**, **Table 2**). To determine if the observed variability is due to the cellular model that we use, or not, we also used two other models: another stable model of TREK-1 overexpression (Andharia *et al*, 2017) and transiently transfected HEK 293T cells with TREK-1 (pIRES2 *KCNK2* WT) (**Figure 3C**). The I_PUFA_/I_0_ variability observed can be explained by the variety of constitutively active TREK-1 channels at resting condition.

**Figure 3:**
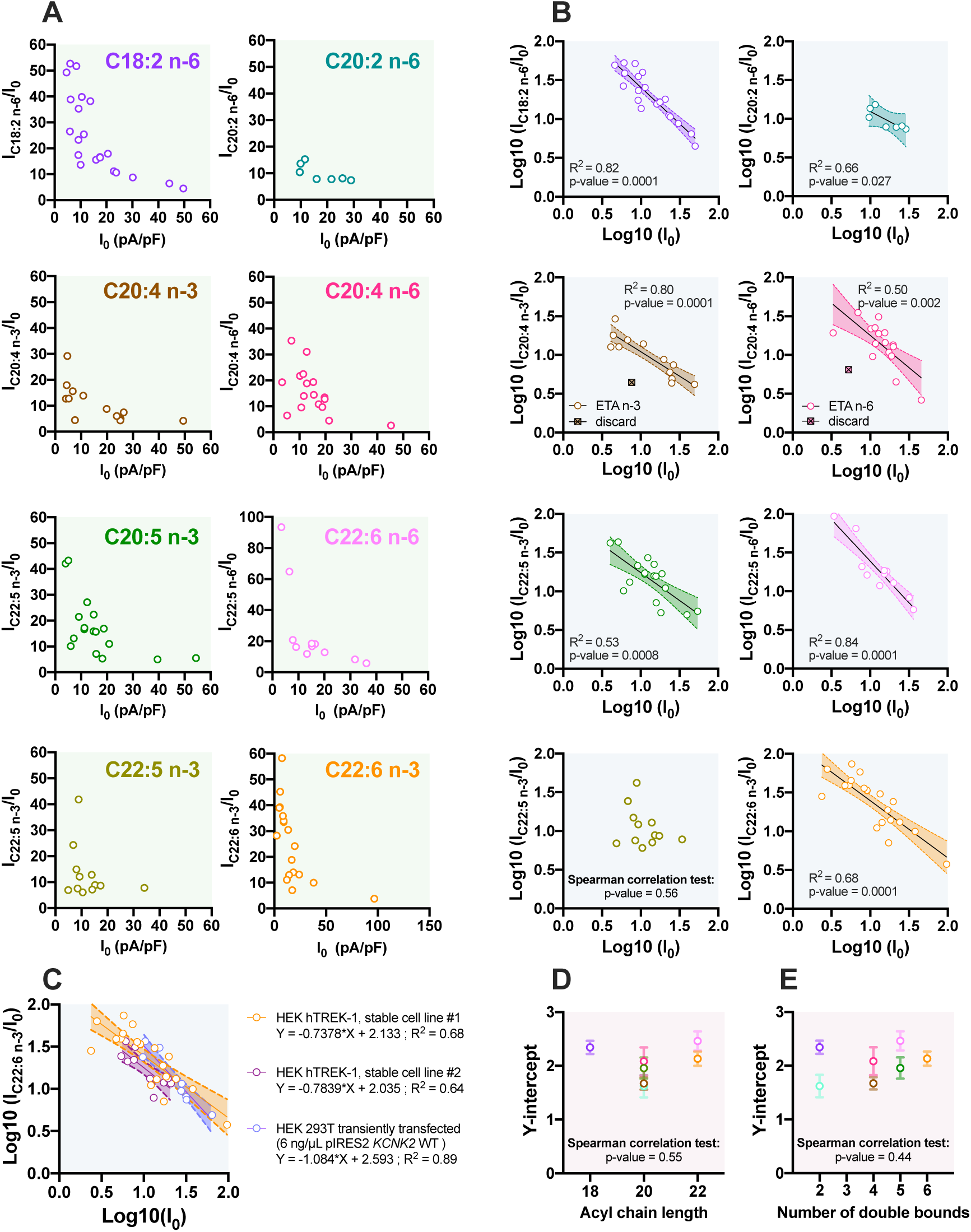
Variability in TREK-1 activation by PUFAs depends on the initial current I_0_. A. Scatter plot representations of I_PUFA_/I_0_ as a function of I_0_. **B.** Linearization of the I_PUFA_/I_0_ *vs* I_0_ relationship following log-transformation for each PUFA. The simple linear regressio () are represented with the 90% confidence interval (IC; dotted line) used to discard one of the C20:4 n-3 and C20:4 n-6 data points (). A point is discarded only if it is out by more than twice the 90% IC. **C.** Relationship between Log10(I_C22:6 n-3_/I_0_) and Log10(I_0_) at 0 mV in different cell lines: HEK hTREK-1 stable cell line #1 (orange), HEK hTREK-1 stable cell line #2 (red) and HEK 293T cells transiently transfected with TREK-1 channel (pIRES2 KCNK2 WT 6 ng/µL)(purple). **D.** Lack of correlation between the Y-intercept of the linear regressions obtained before and the acyl chain length, **E.** and the number of double bounds, for each PUFA at 10 µM.

**Table 2:**
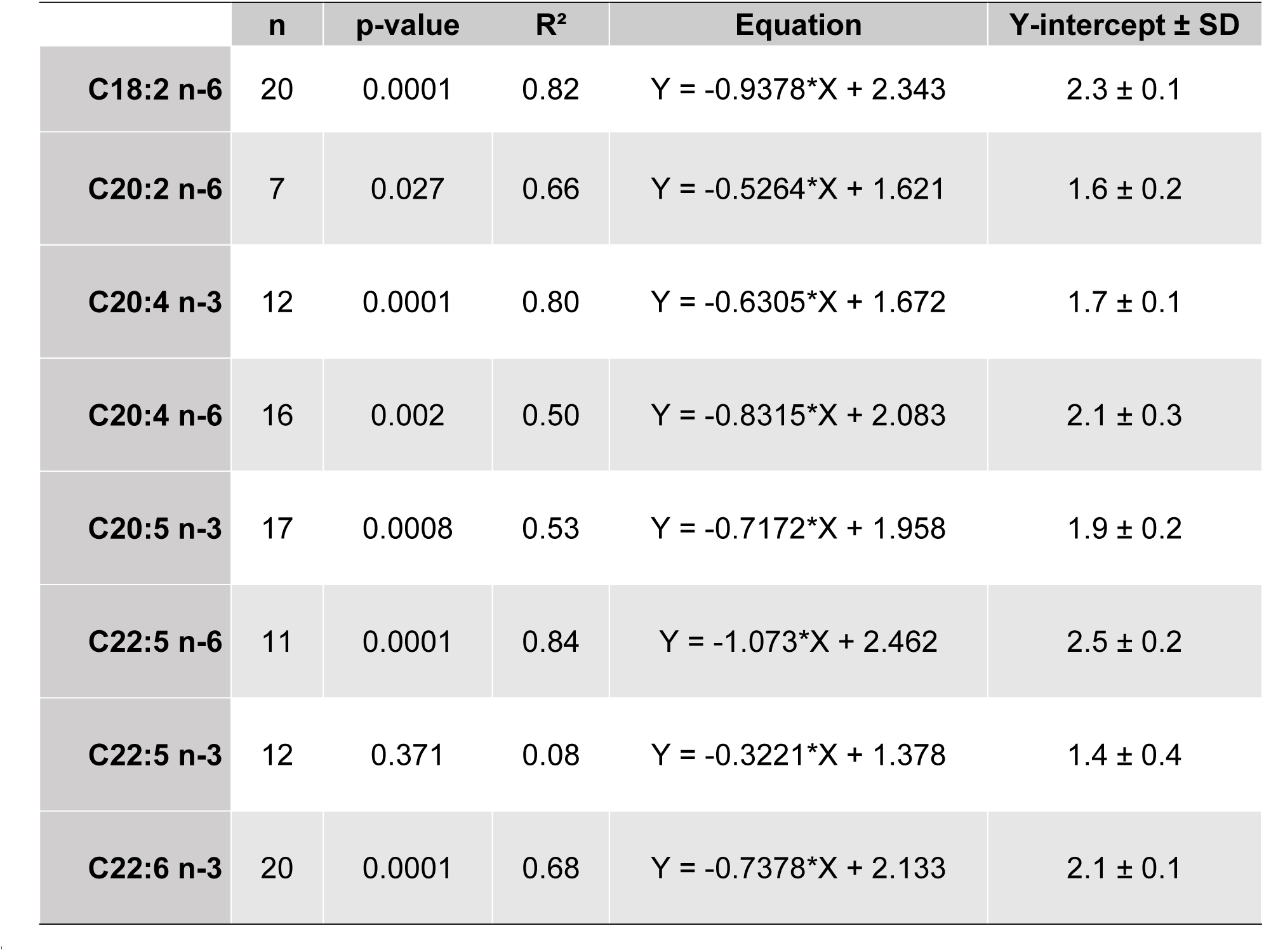
Parameters of the linear regression Log10 (I_PUFA_/I_0_)=f(Log10(I_0_)). TREK-1 current was recorded in whole-celle configuration of patch-clamp.The p-value indicates the significativity of the relationship, the R^2^ indicates the goodness of the fit and the Y-intercept reflects the fold-activation of TREK-1 current for an unitary current as Log(1)=0.

|  | n | p-value | R<sup>2</sup> | Equation | Y-intercept ± SD |
| --- | --- | --- | --- | --- | --- |
| **C18:2 n-6** | 20 | 0.0001 | 0.82 | Y = -0.9378*X + 2.343 | 2.3 ± 0.1 |
| **C20:2 n-6** | 7 | 0.027 | 0.66 | Y = -0.5264*X + 1.621 | 1.6 ± 0.2 |
| **C20:4 n-3** | 12 | 0.0001 | 0.80 | Y = -0.6305*X + 1.672 | 1.7 ± 0.1 |
| **C20:4 n-6** | 16 | 0.002 | 0.50 | Y = -0.8315*X + 2.083 | 2.1 ± 0.3 |
| **C20:5 n-3** | 17 | 0.0008 | 0.53 | Y = -0.7172*X + 1.958 | 1.9 ± 0.2 |
| **C22:5 n-6** | 11 | 0.0001 | 0.84 | Y = -1.073*X + 2.462 | 2.5 ± 0.2 |
| **C22:5 n-3** | 12 | 0.371 | 0.08 | Y = -0.3221*X + 1.378 | 1.4 ± 0.4 |
| **C22:6 n-3** | 20 | 0.0001 | 0.68 | Y = -0.7378*X + 2.133 | 2.1 ± 0.1 |

Thanks to the linear regression analysis, we obtained the Y-intercept which reflects the fold-activation of TREK-1 current for an unitary current (Log(1)=0). The Y-intercept of DPA n-3 cannot be calculated since there was no correlation between I_PUFA_/I_0_ and I_0._ For the 7 others PUFAs, the Y-intercept values allow their separation into 3 groupes with the following activation sequence: C22:6 n-3, C22:5 n-6, C18:2 n-6 > C20:5 n-3, C20:4 n-6 > C20:4 n-3, C20:2 n-6 (**Table 2**). Then, plotting the relationship between Y-intercept (Y axis) and the number of carbons (X axis) reveled once again that there is no correlation between the fold-activation of TREK-1 and the acyl chain length of PUFAs (**Fig 3D**). There is also no correlation with the number of double bounds (**Fig 3E**). In conclusion, TREK-1 activation by PUFAs is definitively not dependent of the acyl chain length and the number of double bond (**Fig 1D-G** and **Fig 3D-E**).

### Variable activation rate suggests different binding affinities of PUFAs for TREK-1 channel

To explore a new way of action of PUFAs on TREK-1 channel, we analyzed and compared the kinetics of activation of TREK-1 perfusing PUFAs or ML402, a binding activator of TREK-1. **Fig 4A to C** represent the mean ± SEM of the normalized current densities (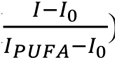) over the time in response to PUFA n-6, PUFA n-3 and ML402, respectively. Representative traces of current densities activation are presented on the right panels. Activation kinetics were fitted with a sigmoid equation (see material and method **equation (3)**) and the activation rate (min^-1^) was calculated as the inverse of the slope of the sigmoid (**Fig 4D**). We were able to distinguish at least two types of activation rate, a fast one above 3 min^-1^ (C18:2 n-6, C22:6 n-3 and ML402) and a slow one less than 3 min^-1^ (C20:2 n-6, C20:4 n-3, C20:4 n-6, C20:5 n-3, C22:5 n-6 and C22:5 n-3). As reported in **Table 3**, the averaged half-activation of TREK-1 channel was smaller for C18:2 n-6 and C22:6 n-3, both having comparable kinetics to those observed for ML402. Although there were no significant differences between these three compounds and C22:5 n-6 and C20:5 n-3, the kinetics of the latters appeared slightly slower (**Fig 4D**, **Table 3**). In contrast, C20:4 n-3, C20:4 n-6 and C22:5 n-3 had slower kinetics with an averaged half-activation close to 4 min (**Table 3**). Since C18:2 n-6 and C22:6 n-3 have the same fast activation rates and C22:5 n-3 has the slowest one, we assumed that the activation rate of the TREK-1 by PUFAs does not depend on the acyl chain length. However, among the PUFAs, there is a positive correlation between the activation rate (min^-1^) and the fold-increase of TREK-1 current (I_PUFA_/I_0_) (Spearman correlation test: p-value = 0.007 and r = 0.88; **Fig 4E**). As the stronger activators are the faster activators of TREK-1, we proposed that some PUFAs, as C18:2 n-6 and C22:6 n-3 have a higher binding affinity for TREK-1 which would allow them to activate it faster and stronger.

**Figure 4:**
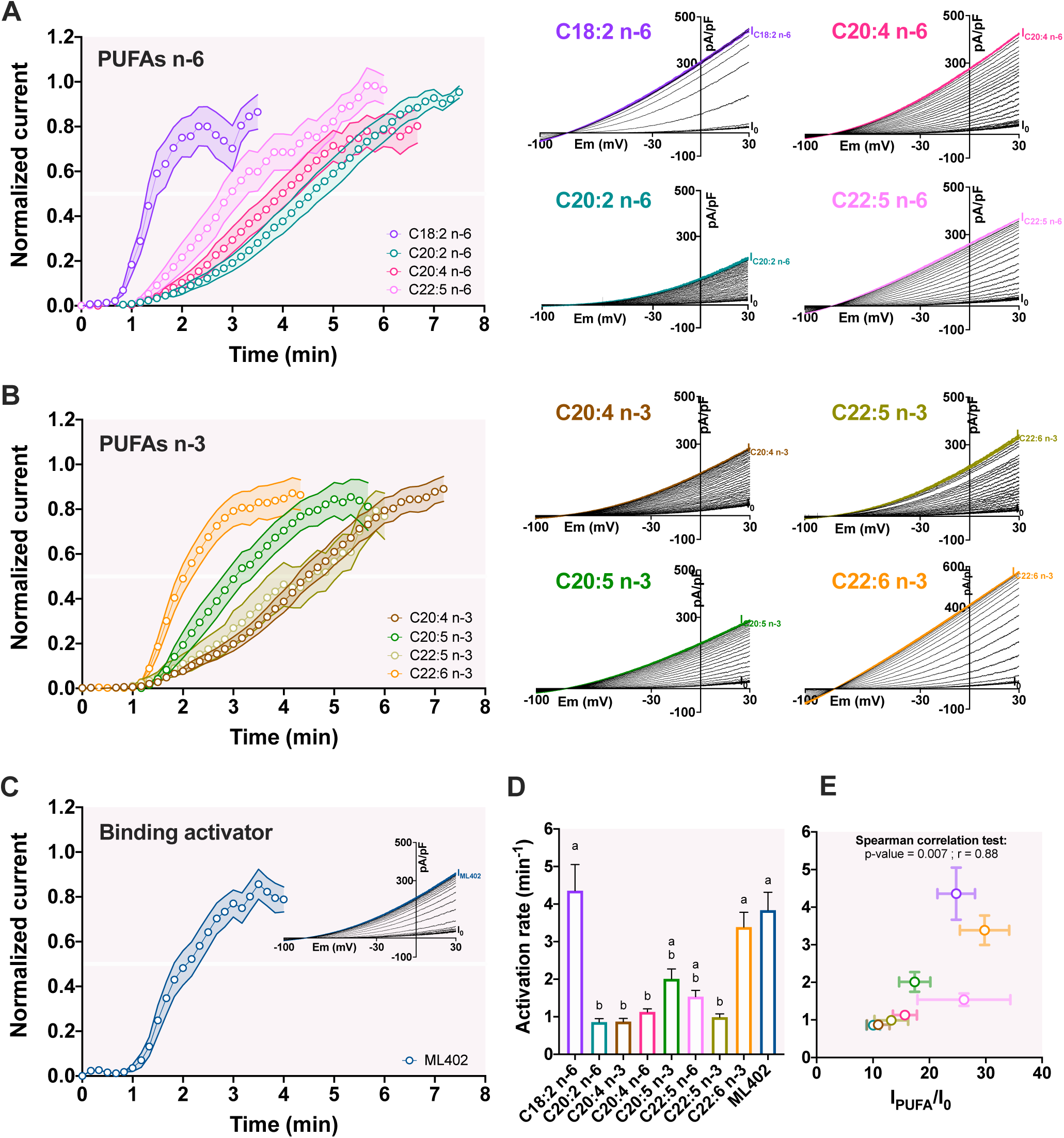
Activation kinetics of TREK-1 channel by PUFAs. A-C. Time course showing the effects of 10 µM PUFAs and ML402 on I_TREK-1_ at 0 mV. PUFAs and ML402 were superfused until the steady-state was reached. *Insets* show the representative current densities recorded in control medium and then under PUFAs or ML402 application. **D.** Bar graphs of the activation rate **+** SEM to reach the steady-state of I_PUFA_. Groups were compared with a Kruskal-Wallis test followed with the *post-hoc* Dunn’s test. Two bars having the same letter are not significantly different. **E.** Relationship between the fold-increase of TREK-1 (I_PUFA_/I_0_) and the activation rate (min^-1^) (R^2^=0.60; p-value=0.02; error barres show the SEM of the I_PUFA_/I_0_ and the activation rate).

**Table 3:** Parameters of the activation kinetic of TREK-1 by PUFAs and ML402 in whole-cell configuration of patch-clamp. Values indicate the time (min) needed to reach the half-activation and the steady-state of the activation of TREK-1. The activation rate (min^-1^) derived from the sigmoidal fit as: 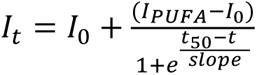 and activation rate 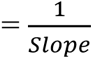.

|  |  | Half-activation (min) | Steady-state (min) | Activation rate (min<sup>-1</sup>) |
| --- | --- | --- | --- | --- |
|  | n | Mean ± SEM | Mean ± SEM | Mean ± SEM |
| **C18:2 n-6** | 20 | 1.9 ± 0.2 | 3.7 ± 0.3 | 4.5 ± 0.7 |
| **C20:2 n-6** | 7 | 4.6 ± 0.3 | 7.4 ± 0.3 | 0.9 ± 0.1 |
| **C20:4 n-3** | 12 | 4.6 ± 0.2 | 7.1 ± 0.2 | 0.9 ± 0.1 |
| **C20:4 n-6** | 16 | 3.9 ± 0.3 | 6.7 ± 0.3 | 1.1 ± 0.1 |
| **C20:5 n-3** | 17 | 3.2 ± 0.3 | 5.7 ± 0.3 | 2.0 ± 0.3 |
| **C22:5 n-6** | 11 | 3.1 ± 0.3 | 5.3 ± 0.4 | 1.5 ± 0.2 |
| **C22:5 n-3** | 12 | 4.2 ± 0.4 | 5.9 ± 0.4 | 0.9 ± 0.1 |
| **C22:6 n-3** | 20 | 2.3 ± 0.2 | 4.3 ± 0.3 | 3.4 ± 0.4 |
| **ML402** | 19 | 2.1 ± 0.2 | 3.9 ± 0.3 | 3.8 ± 0.5 |

### Activation of TREK-1 channel by PUFAs is fully reversible

To see if the activation of TREK-1 channel by PUFA is due to an insertion and thus a modification of membrane tension, we looked at the washout kinetic with extracellular medium free of Bovin Serum Albumin (BSA). We focused on C18:2 n-6, C20:5 n-3 and C22:6 n-3, the most potent activators of C18, C20 and C22 PUFAs, respectively (**Table 1**). ML402 activation reversed immediately and 50% of washout occurred in less than 1 min (**Fig 5A and 5B**). C20:5 n-3, had a kinetic of washing (washout 50%: 0.9 ± 0.1 min) comparable to ML402 (**Fig 5A and 5B**). Even though the washout of C18:2 n-6 and C22:6 n-3 was slower than ML402 (washout 50%, mean ± SEM: 2.4 ± 0.2 min, 3.7 ± 0.3 min and 0.4 ± 0.04 min, respectively), PUFAs effect were also fully reversed under washing. Once again, there is no correlation between the acyl chain length (**Fig 5C**, Spearman correlation tes: p-value > 0.99) or the number of double bounds (**Fig 5D**, Spearman correlation test: p-value > 0.99) and the time needed to reverse TREK-1 activation. C18:2 n-6 and C22:6 n-3, that activated TREK-1 at least twice more than ML402 (I/I_0_: 24.8 ± 3.3, 29.8 ± 4.4 and 9.6 ± 0.9, respectively), had a total reversibility in few minutes. At this point, we cannot exclude a membrane insertion of PUFAs, but we assume that the main effects of PUFAs on TREK-1 activation could be a direct and reversible interaction of PUFAs with the channel, as ML402, or as it is well known for KCNQ1 (Liin *et al*, 2015) and the Shaker H4 Kv channel (Börjesson *et al*, 2008)

**Figure 5:**
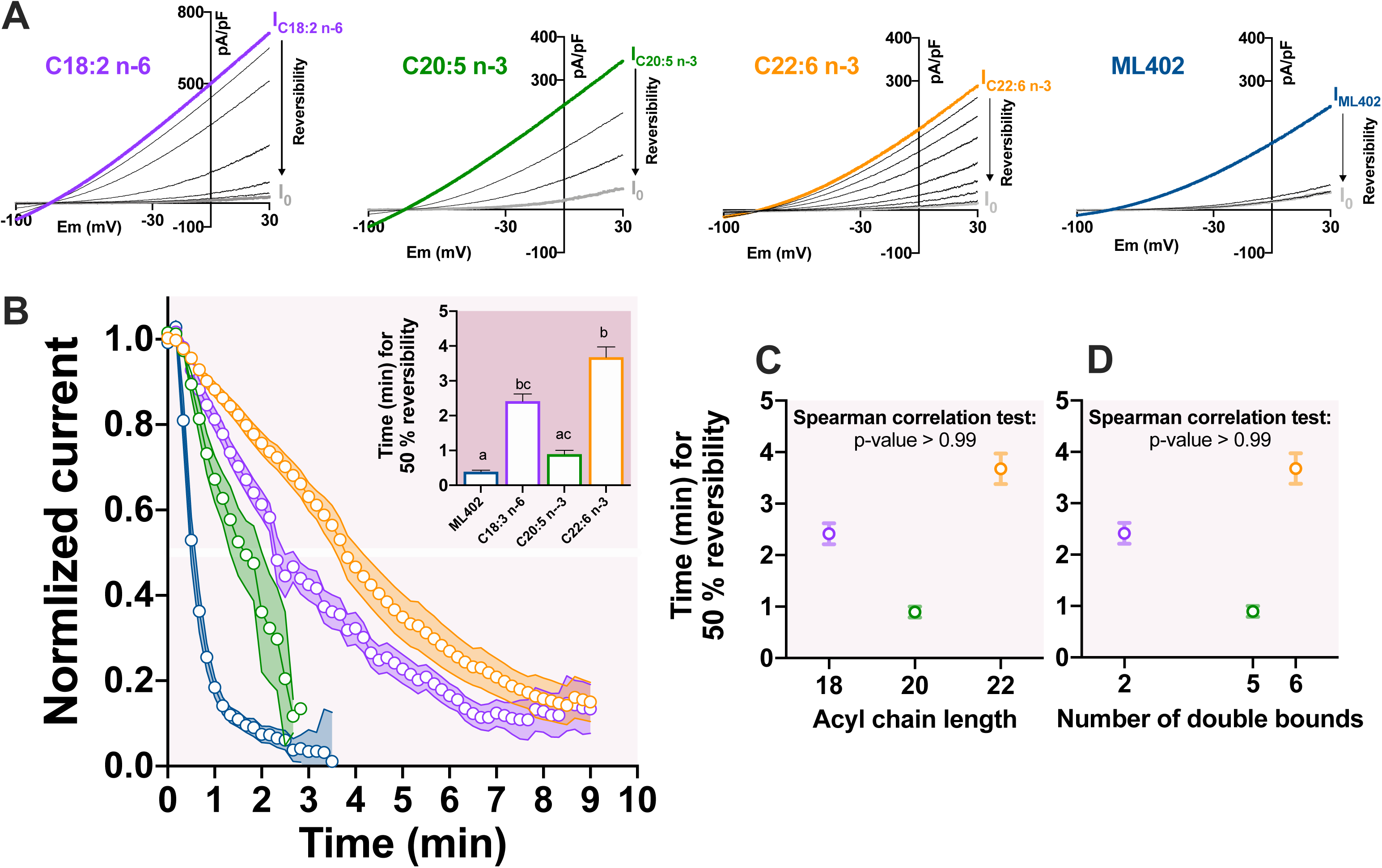
Washout kinetics of LA, EPA, DHA and ML402. A. Representative traces of the reversibility of the current density with one trace per minute (grey: initial current I_0_; colors: PUFA-activated current density at steady-state; black: washout current density). **B.** Normalized current-time curve of the reversibility for each compound (mean ± SEM) with 1 point every 10s. The current was normalized as (I-I_0_)/(I_PUFA_-I_0_). The inset shows the time to reach 50% of the reversibility of TREK-1 activation. **C.** Lack of correlation between the time for 50 % of washout and the acyl chain length and **D.** the number of double bounds.

### Alteration of membrane curvature or fluidity did not explain activation of TREK-1 channel by PUFAs

In order to evaluate the membrane curvature and tension effects on the PUFAs-induced TREK-1 activation, we performed experiments in the inside-out configuration of the patch-clamp technique (+30 mV, symmetrical condition: 145 mM KCl). In this configuration, we were able to superfuse molecules at the inner face of the membrane and PUFAs must induce a curvature of the membrane opposite to the one obtained in the whole-cell configuration. As shown in **Figure 6A**, ML402 superfusion at the inner face of the membrane induced a reversible increase of TREK-1 current (**Table 4**). Kinetics of activation and washout were comparable to those obtained in the whole-cell configuration (**Fig 4** and **Fig 5**) suggesting that the ML402-binding site is accessible from the outer and the inner leaflet of the membrane. Interestingly, a comparable reversible activation of TREK-1 channel was obtained for C18:2 n-6 and C22:6 n-3 5 µM, suggesting that the membrane curvature is not involved in the activation of TREK-1 channel by PUFAs (**Fig 6B** and **6C**, **Table 4**).

**Figure 6:**
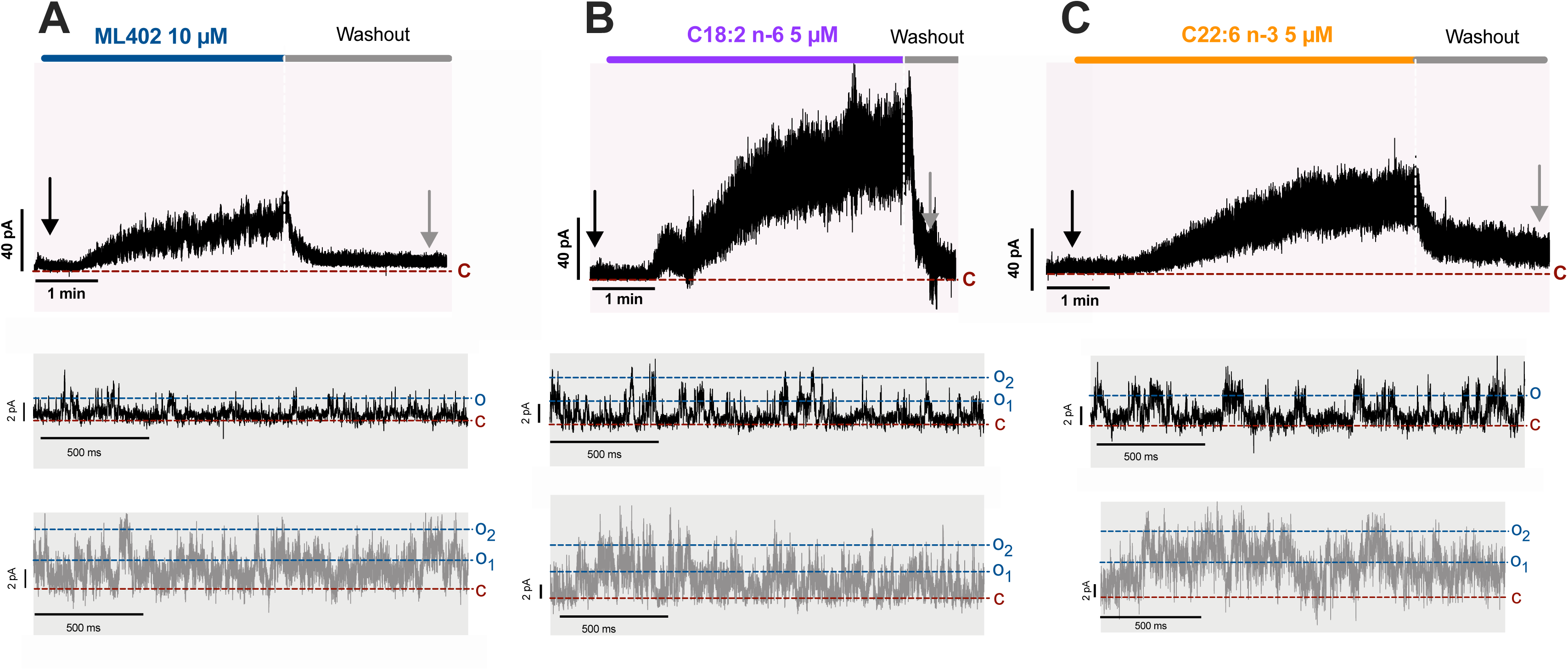
PUFAs activate TREK-1 in inside-out configuration of patch-clamp. A-C. Representative traces showing the activation and reversibility in inside-out configuration of patch-clamp technique for **A.** ML402 10 µM **B.** C18:2 n-6 5 µM and **C.** C22:6 n-3 5 µM superfused at the inner face of the membrane. Membrane potential was held at +30 mV. The expanded current traces where extracted at the time indicated by the arrows.

**Table 4:**
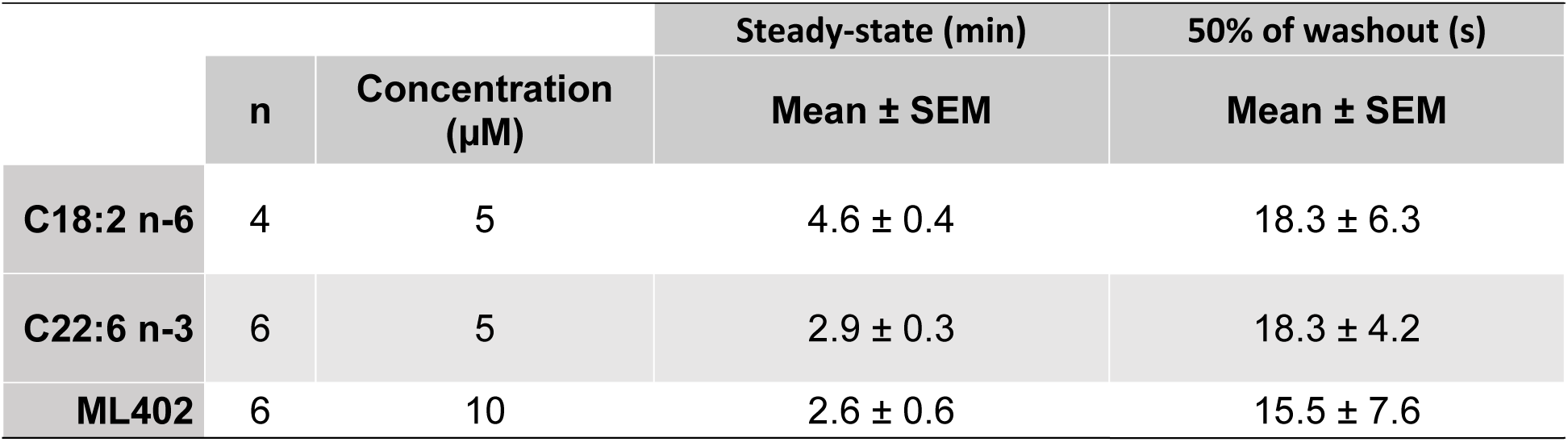
Descriptive statistics for TREK-1 activation and washout kinetics in inside-out configuration of patch-clamp. Values indicate the time needed to reach the steady-state (min) and the time to reach 50% of washout (s).

|  |  |  | Steady-state (min) | 50% of washout (s) |
| --- | --- | --- | --- | --- |
|  | n | Concentration (μM) | Mean ± SEM | Mean ± SEM |
| **C18:2 n-6** | 4 | 5 | 4.6 ± 0.4 | 18.3 ± 6.3 |
| **C22:6 n-3** | 6 | 5 | 2.9 ± 0.3 | 18.3 ± 4.2 |
| **ML402** | 6 | 10 | 2.6 ± 0.6 | 15.5 ± 7.6 |

Then, we assessed the membrane fluidity changes during PUFAs application with a pyrenedecanoic acid probe (PDA), analog to lipids. By measuring the ratio of PDA monomer to excimer fluorescence (405nm/470nm ratio), a quantitative assesment of the membrane fluidity can be obtained at different time points by following the ratio modification over time (F/F_0_-1). We focus our experiments on C18:2 n-6 and C22:6 n-3 which are the stronger activators of TREK-1 and C18:3 n-3 that failed to activate TREK-1. These 3 PUFAs at 10 µM did not modify the membrane fluidity even after 50 minutes of application, while at 100 µM they induced a decreased of F/F_0_-1 from T_0_ and compared to the control condition (basic extracellular medium). These results indicate that membrane fluidity is not modified by PUFAs at 10 µM (**Fig 7A-C**), at least within 50 minutes of application. In addition, given that TREK-1 activation starts at 1 minute of perfusion of C18:2 n-6 and C22:6 n-3 10 µM (**Fig 4**) it is unlikely that PUFA effects on TREK-1 activation are due to an increase in membrane fluidity. Altogether, these data suggest that at least both C18:2 n-6 and C22:6 n-3 PUFAs activate TREK-1 channel by direct interaction with TREK-1 protein and not by a modification of the membrane fluidity.

**Figure 7:**
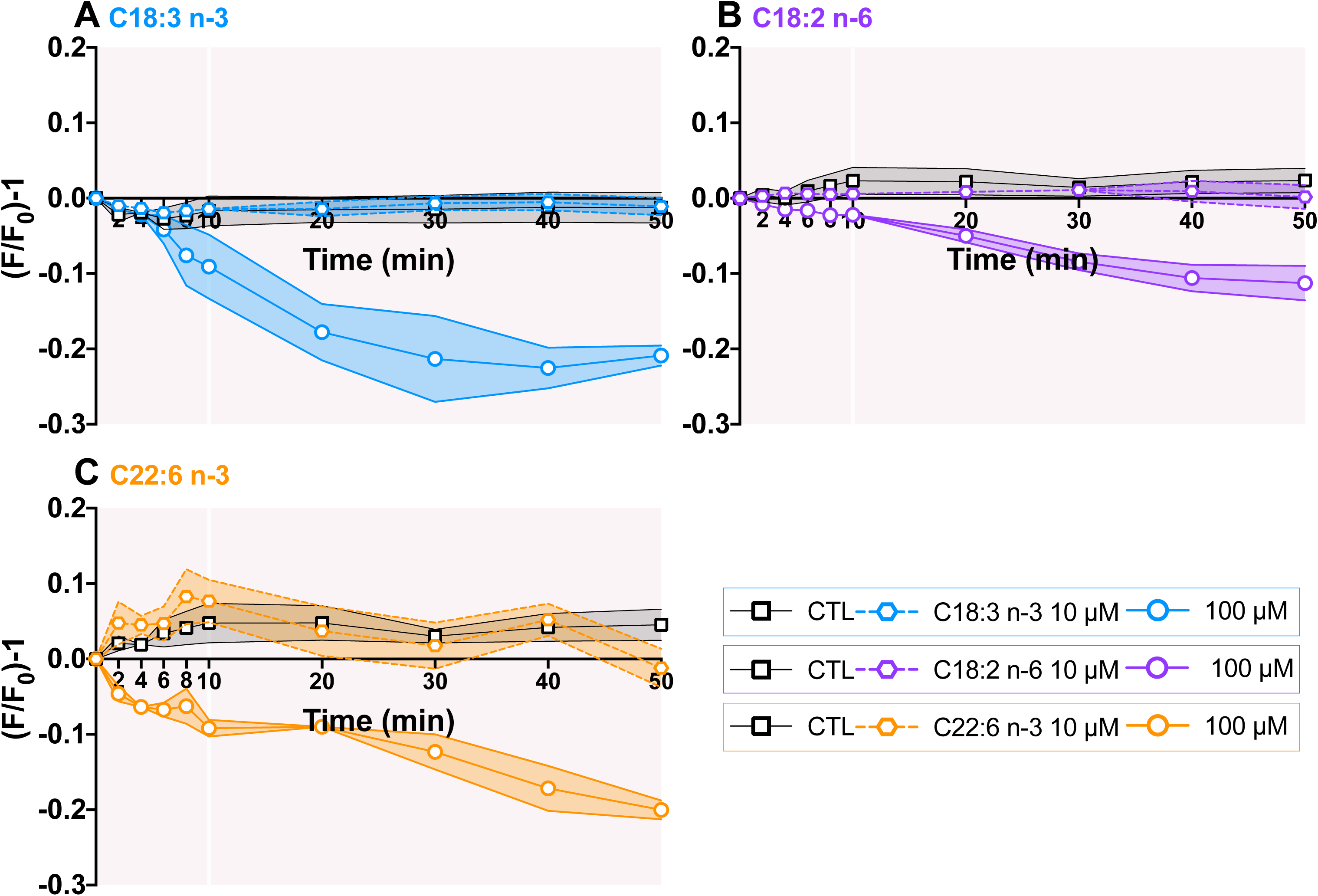
Membrane fluidity is not altered by PUFAs at 10 µM A-C. Membrane fluidity is not changed over time by 10 µM of PUFA but 100 µM:**A.** C18:3 n-3 (10 µM: n=3; 100 µM n=6; CTL n=4), **B.** 18:2 n-6 (10 µM: n=6; 100 µM: n=7; CTL n=6), **C.** C22:6 n-3 (10 µM: n=5; 100 µM: n=5; CTL n=5),

### DHA interacts directly with TREK-1 channel protein in TREK-1 enriched microsomes

In order to assess a potential direct PUFA-TREK-1 interaction, we purified microsomes from hTREK-1/ HEK and native HEK 293T cells and labeled lysine residues of the total proteins to perform affinity measurements using Spectral Shift (SpS). At first glance, we observed similar affinity from C22:6 n-3 in these two type of microsomes: K_d,TREK-1_ ∼ 50 µM and K_d,HEK_ ∼ 100 µM highlighting a similar mode of association of C22:6 n-3 within the microsomes. However, the SpS signal displayed subtle differences according to whether or not microsomes were enriched in TREK-1 protein. Knowing SpS signal rises from fluorescence recorded by two individual channel, and having a strong reproducibility from each condition, we can derive the following assumption:

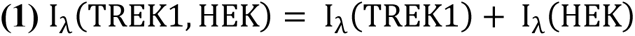

Where for each individual wavelength (λ), the fluorescence recorded for the TREK-1-enriched microscomes correspond to both the fluorescence from the labelled empty microsomes and from the labelled TREK-1.

Being able to isolate the specific fluorescence associated to TREK-1 *(I_λ_(TREK1))* within the TREK-1-enriched microsomes (*I_λ_(TREK1, HEK))*, we can use individual channel for further calculation using **equation 2**, isolating the SpS signal associated only to TREK-1 by normalizing out the background signal coming for the free microsomes.

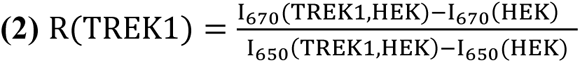

When performing so, the specific TREK-1 dose response to C22:6 n-3 gives a K_d_ = 44 µM (**Fig 8**). Despite displaying similar affinities, the TREK-1-enriched microsomes shows a statistically better affinity than the microsomes themselves. However, it is worth noticing only a 2-fold increase of affinity, which may highlight a similar binding mode for both interactions. Altogether, this suggests an interaction mediated by the lipid bilayer, such as a membrane insertion followed by interaction with TREK-1 channel.

**Figure 8.**
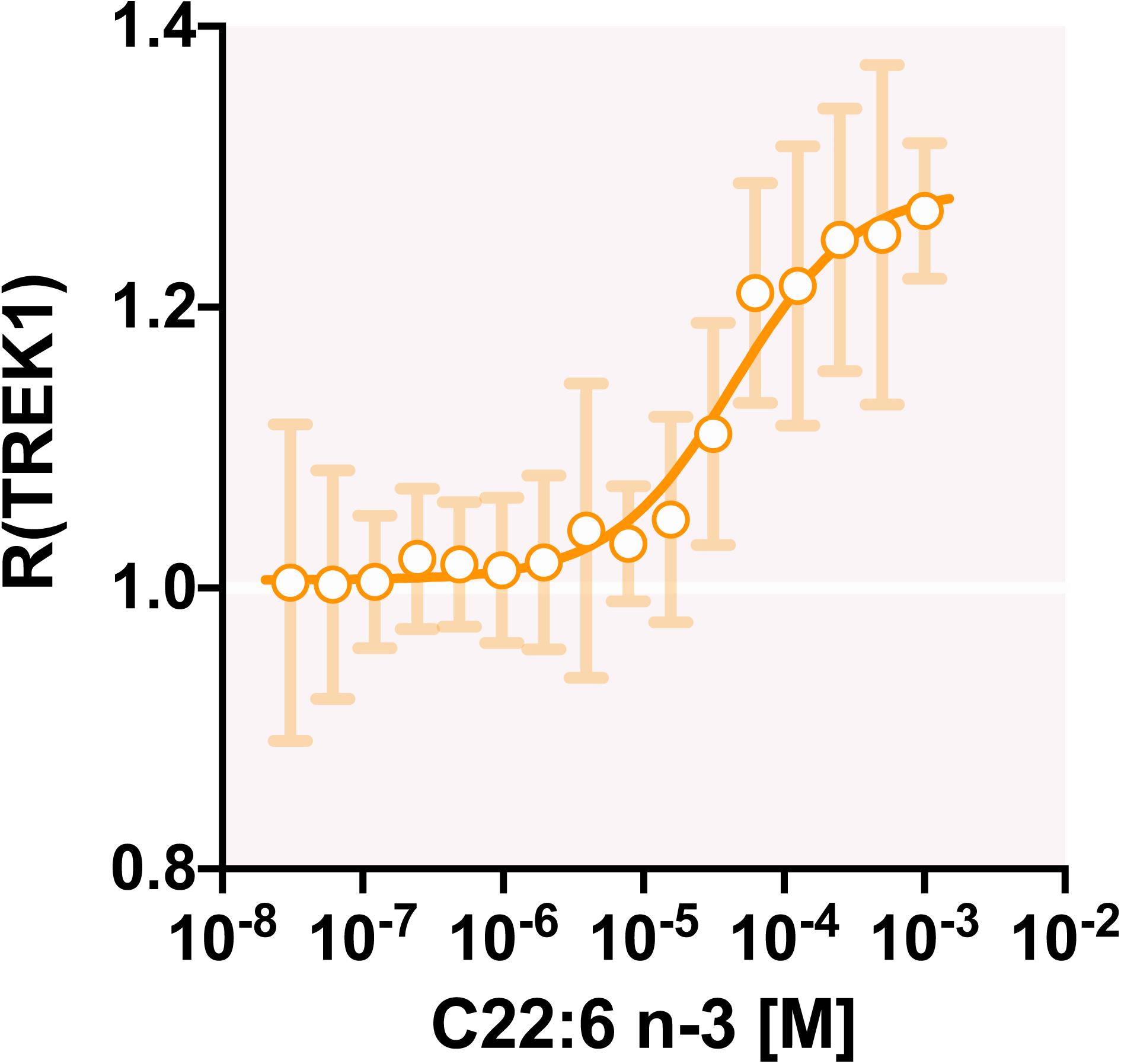
: Effect C22:6 n-3 on SpS signal in TREK-1-enriched microsomes and non-enriched microsomes. (*I_λ_(TREK1, HEK)* represents the binding affinity of C22:6 n-3 (from 1 mM to 30.5 nM) for TREK-1-enriched microsomes from 1 mM to 30.5 nM (n=5-9 experiments).

## DISCUSSION

### TREK-1 channel activation by PUFAs does not involve mechano-sensitivity but a direct interaction

In this study, we reported that TREK-1 channel is reversibly activated by polyunsaturated fatty acids (PUFAs), as already shown in different studies, each being focused mainly on one PUFA: AA (Patel *et al*, 1998); LA (Danthi *et al*, 2003); DHA (Ma & Lewis, 2020). Our study is the first to compare the effects of different PUFAs having between 18 to 22 carbon atoms and 2 to 6 double bonds on TREK-1 channel. We demonstrate that C22:6 n-3 and C18:2 n-6 are the most potent activators of TREK-1 with an activation as fast as the one of the direct activator ML402 and a fully reversibility.

TREK subfamily of K2P channels that includes TREK-1, TREK-2 and TRAAK, is characterized by a mechano-sensitivity and therefore, channels could feel changes in the membrane curvature induced by PUFAs insertion as for CPZ (Patel et al., 1998). As PUFAs are anionic amphipath compounds, with a hydrophilic carboxyl group and a lipophilic tail, they preferentially insert into the outer leaflet of the membrane which is positively charged (Martinac *et al*, 1990; Sheetz & Singer, 1974). Thus, the longer the lipophilic carbon chain is, the more the PUFAs will be inserted into the membrane, modifying the local membrane elastic properties (curvature and fluidity) (Leifert *et al*, 1999). Also, for a given carbon chain length, the fluidity increases with the number of double bonds. According to the bilayer-couple hypothesis, PUFA effects on TREK subfamily channels were supposed to be due to a modification of elastic properties of the membrane leading to an increase of the tension transmitted to the channels (Patel *et al*, 1998). The study of the TRAAK channel activation by PUFAs (C18:2 n-6, C20:4 n-6, C20:5 n-3, C22:6 n-3) in the excised patch configuration shows that TRAAK activation is positively correlated with the carbon chain length of PUFAs and the number of double bonds (Fink *et al*, 1998; Patel *et al*, 2001). However, we found no correlation between the acyl chain length, the number of double bonds and the potentiation of I_TREK-1_. In the opposite, C18:2 n-6 and C22:6 n-3, respectively the shortest and the longest PUFA tested, are the most potent activators of TREK-1 channel. It is worth to note that C18:3 n-3, which differs only by one double bond from C18:2 n-6, failed to activate TREK-1 channel. Also, C22:6 n-3 having the same number of carbons than C22:5 n-3 is more than twice as effective in activating TREK-1. PUFAs having intermediate acyl chain length produce intermediate activation of TREK-1, in the same range as the direct activator ML402, independently of the double bonds number. In that respect, the bilayer-couple hypothesis suggesting an activation of TREK-1 by PUFAs-induced mechanosensitive pathway is not appropriate. Similar results were obtained in the literature on TREK-2 channel study, C20:4 n-6 being less efficient than C22:6 n-3 and C18:2 n-6 (Lesage *et al*, 2000).

To better characterize the mechanism of action of C22:6 n-3 and C18:2 n-6 on TREK-1, we compared their kinetics of activation and reversibility with those of ML402. ML402 is a direct activator of TREK-1, binding within a cryptic pocket behind the selectivity filter that directly stabilize the C-type gate (Lolicato *et al*, 2017). The activation kinetic of TREK-1 by C22:6 n-3, C18:2 n-6 and ML402 are comparable and faster than the other PUFAs. This suggests a possible interaction of C22:6 n-3 and C18:2 n-6 with the channel like the activator ML402 (Lolicato *et al*, 2017) and the inhibitor norfluoxetine (Dong *et al*, 2015). This hypothesis is reinforced by the inside-out experiments where C22:6 n-3, C18:2 n-6 and ML402 were applied on the inner leaflet of the membrane. Although the PUFA insertion must induce opposite curvartures while they insert from the inner (inside-out configuration) or the outer leaflet (whole-cell configuration), they still activate TREK-1 channel. We finally propose that at least C22:6 n-3 and C18:2 n-6, like ML402, interact with the channel on an accessible site from both the inner and the outer leaflet of the cell, and suggesting a binding site accessible via the lipid bilayer. Studies have already hypothesized that arachidonic acid (C20:4 n-6) could act directly by interacting with the channel (Maingret *et al*, 1999, 2000) and others have already shown that free PUFAs can directly interact with ionic channels. Indeed, C22:6 n-3 and C18:3 n-6 interact with K_v_7.1 (*KCNKQ1*)(Liin *et al*, 2015; Yazdi *et al*, 2021). PUFAs also interact with Shaker H4 Kv channel closed to the voltage-sensor domaine through the negatively charged carboxyl group (Börjesson *et al*, 2010; Börjesson & Elinder, 2011; Börjesson *et al*, 2008). As TREK-1 lacks a canonical voltage-sensor domain, we can hypothesize that there is another lipophilic binding site in TREK-1 interacting with the carboxyl head of PUFAs. This hypothetical PUFA binding site on TREK-1 does not correspond to the binding site of ML402 since its mutation does not prevent activation by C20:4 n-6 (Lolicato et al., 2017). Moreover, since the steady-state of TREK-1 activation by PUFA and ML402 was reached in a range of the minutes, we suggest that the PUFAs binding-sites have a limited access (Maingret *et al*, 2000). It has to be noted that bovine serum albumin (BSA) is not required to get a reversal effect during the washout as it is supposed to be when the effects are due to membrane insertion of the PUFAs (Kang & Leaf, 1994; Leifert *et al*, 1999).

Finally, in both whole-cell and inside-out configurations, the washout kinetics of PUFAs were immediate and the initial current recovered in few minutes. Despite a total reversibility of the TREK-1 activation under washout, we cannot definitively exclude a membrane insertion of PUFAs owing to their lipophilic properties. Nevertheless, if PUFAs insert into the membrane, they do not modify the biophysic properties of the membrane (fluidity, curvature, tension) at 10 µM over a 50 min periode whereas TREK-1 is fully activated by PUFAs in few minutes. Affinity measurements between PUFAs and TREK-1-enriched microsomes effectively indicates a binding with the lipid bilayer as observed for the empty microsomes. However, once the signal due to PUFA interaction with the bilayer is subtracted from the total signal, a direct binding of PUFAs to TREK-1 is measurable. This specific interaction displays a stronger affinity (K_D,TREK-1_ *∼* 44 µM) than the simple signal involving PUFA binding with the bilayer of the microsomes. Therefore, we propose that two mechanisms act together to increase I_TREK-1_: (1)PUFA insertion into the membrane, with no modification of its elastic properties, is probably recquired to reach (2) a lipophilic binding site on TREK-1 channel accessible via the lipid bilayer as for the Shaker channel (Börjesson & Elinder, 2011)

### Initial TREK-1 variability influence PUFAs response

An unexpected result of this study is the observation that the PUFA effects depend on the TREK-1 initial current. This variability in initial I_TREK-1_ might have many origins such as different levels of post-translationnal modifications, different channels recruitment into the membrane or even the presence of 2 different conductances of TREK-1 channel at the single channel level (Xian Tao Li *et al*, 2006; Andharia *et al*, 2017). A comparable variability in the current-fold increase induced by PUFAs was already observed for TREK-1 activation by AA (Maingret *et al*, 2000), TRAAK (Fink *et al*, 1998; Patel *et al*, 2001) and TREK-2 (Bang *et al*, 2000; Lesage *et al*, 2000) but never explained.

It is important to note that all these studies were performed on COS-7 cells or HEK-293 cells where mammalian post-translational modifications exist. In the study of Ma and Lewis in 2020, whole-cell recording of TREK-1 and TREK-2 by arachidonic acid were performed in oocytes (*Xenopus Laevis*) and no such variability in the current fold-increase was observed. It is known that in this heterologous expression model, post-translational modification are different (Mantegazza *et al*, 2010). Therefore, the possibility that constitutive TREK-1 current varies according to the phosphorylation level (or other modifications) and influence the effects of PUFAs should be taken in consideration. It is known that TREK-1 channel is modulated by intracellular pathways, related to PKA, PKC and PKG signalizations (Koh *et al*, 2001; Murbartián *et al*, 2005; Honoré, 2007), but to the best of our knowledge, no study between PUFA activation of TREK-1 and phosphorylation levels has been performed so far. As a consequence of this variability, to study the effect of an activator of I_TREK-1_, we need a sufficient number of cells to apply the Y-intercept of the regression line, probably the best indicator of the degree of activation of TREK-1. The existence of such variability should be taken into consideration in pathophysiological studies associated with variability of expression or response, potentially demultiplying the heterogeneity of TREK-1 response to activators.

Surprisingly, one of the most efficient activator of TREK-1 channel is C18:2 n-6. As TREK-1 is expressed in cardiomyocytes (Li & Toyoda, 2015; Kelly *et al*, 2006; Decher *et al*, 2017) C18:2 n-6 should modulate action potential shape and resting membrane potential.This suggests that C18:2 n-6 could display potent cardio as well as neuroprotective effects. If anti-arrhythmic properties of omega-3 PUFAs have been thoroughly studied (Kang & Leaf, 1994, 1996; Siscovick *et al*, 2017), the potential anti-arrhythmic effect of omega-6 PUFAs such as C18:2 n-6 has never been considered. C18:2 n-6 can be found in vegetarian and Mediterranean diets, both diets having cardioprotective vertues (de Lorgeril *et al*, 1994; Barnard *et al*, 2019). In both cases the beneficial effects are attributed to the presence of anti-oxidant molecules and C18:3 n-3 but a possible beneficial effect of C18:2 n-6 has never been explored.

## CONCLUSION

To conclude, there is no relationship between the carbon number of the acyl chain, the number of double bonds and the activation of TREK-1 channel. Its most potent activators are C18:2 n-6 (linoleic acid) and C22:6 n-3 (docosahexaenoic acid) and kinetics analysis suggest a direct interaction of PUFAs on TREK-1 through a lipophilic binding site. This direct activation of PUFAs in TREK-1 could require the membrane insertion of PUFAs to facilitate the access to the binding site in the channel.

## MATERIEL AND METHODS

### Cell culture

We used a HEK-293T cell line for spectral shift mesurement and two HEK/hTREK-1 cell lines that stably overexpress the human TREK-1 channel subunit (Moha ou Maati *et al*, 2011; Andharia *et al*, 2017) for electrophysiological experiments and spectral shift mesurement. Cells were grown in an atmosphere of 95% air/5% CO2 in Dulbecco’s modified Eagle’s medium and Glutamax (Invitrogen, Cergy-Pontoise, France) supplemented with 10% (v/v) heat inactivated fetal bovine serum and 0.5 mg/mL G418 to maintain a selection pressure in the HEK/hTREK-1 cell line.

The transient expression of TREK-1 was performed in the HEK 293T cell line. The pIRES2 plasmid in which the coding sequence of TREK-1 (pIRES2 *KCNK2* WT) was inserted, was transfected in HEK 293T cells using the jetPEI® kit (Ozyme) and following the manufacturer protocol. Briefly, 48h before electrophysiological experiments, cells were transfected with a mix of the water-soluble polymer jetPEI® with DNA at 6 ng/mL and then seeded in 35×10 mm dishes (Falcon) in presence of 0.5 mg/mL of G418

### Electrophysiology

Currents were recorded from hTREK-1/HEK-cells using the patch-clamp technique in whole-cell recording (WCR) configuration and in inside-out (IO) configuration. Patch pipette having 2.5-4 MΩ resistances (WCR) and 5-7 MΩ resistances (IO) were obtained from borosilicate glass capillaries by using a two-stage vertical puller (PC-10, Narishige, London, UK). Current acquisition was performed with an Axopatch 200B amplifier (Axon Instrument, Sunnyvale, CA, USA) and low-pass filtered at 5 kHz (WCR) and 2 kHz (IO). Data were digitalized with a digidata 1550B (Axon Instrument, Sunnyvale, CA, USA) at 10 kHz. The pClamp10.7 (Axon Instrument) software was used to impose stimulations protocols and record TREK-1 current. In WCR, cells were kept for experiments if the series resistance were lower than 8 MΩ, the membrane capacitance between 20 and 35 pF.

The experiments were performed at room temperature (∼22 °C). Cells were continuously superfused with a extracellular medium and modulatory compounds at a rate of 1-1.5 mL/min. The dishes volume was kept constant at 2.5-3 mL using a vacuum system, connected to a peristaltic pump (ISMATEC). Using different extracellular potassium concentrations (5 mM KCl or 15 mM KCl), changes in membrane potential were measured in current-clamp allowing to determine the time to change completely the medium around the patched-cells (Le Guennec & Noble, 1994). A complete change of the extracellular medium around the patched-cell occurred in less than 15 seconds.

### Whole-cell recording

the extracellular medium consisted of (in mM): 150 NaCl, 5 KCl, 3 MgCl_2_, 1 CaCl_2_ and 10 HEPES pH adjusted to 7.4 with NaOH. The pipette solution consisted of (in mM): 155 KCl, 3 MgCl_2_, 5 EGTA and 10 HEPES, pH adjusted to 7.2 with KOH. Cells were clamped at a holding potential of −80 mV and superfused with the extracellular medium for over 1 min before the initial current (I_0_) recording. Cells were hyperpolarized to −100 mV for 50 ms and then the macroscopic outward TREK-1 current was elicited with an 800 ms voltage ramp protocol from −100 mV to +30 mV every ten seconds (see the ramp protocole in **Fig 1A**). Thus, the evolution of the current amplitude changes during compound superfusion was followed in real time (**Fig 1B**). A minimum of 3 min superfusion was applied even when the tested compound showed no effect and the superfusion condition was switched once a steady-state was reached. In the case where the tested compound had no effect, a positive control of the TREK-1 activation was performed with 10 µM docosahexaenoic acid (DHA, C22:6 n-3).

### Data analysis (WCR)

Once at steady-state, the current amplitude at 0 mV during the voltage ramp was measured from the average of the last 3 sweeps over a delta of 1 mV (−0,5 mV to 0,5 mV) to get out of the noise. Current amplitudes are expressed in current densities (pA/pF) to reduce the variability due to cell size. Activation an washout kinetics were followed at 0 mV. The activation kinetics from each cell of each compound (PUFAs and ML402) was fitted with a sigmoidal equation (**Fig 5**):

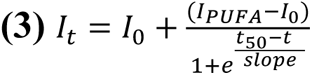

where I_0_ is the initial current and I_PUFA_ is the current caused by PUFA or ML402 at 10 µM concentration. The activation rate (min^-1^) was then calculated as: activation rate 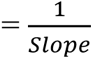.

### Inside-out configuration

bath medium contained (in mM): 140 NaCl, 4.8 KCl, 1.2 MgCl_2_, 10 glucose, 10 HEPES, pH adjusted to 7.4 with NaOH. After excising the membrane patch, the bath medium was replaced by the a medium identical to the pipette medium consisted of (mM): 145 KCl, 1.2 MgCl_2_, 10 glucose, 10 HEPES, pH adjusted to 7.2 with KOH. Cells were clamped at a holding potential of 0 mV, the theorical equilibrium potential for K^+^ ions in this condition. Then, the amplitude current was followed at +30 mV in real time during the perfusion of compounds and the washout.

### Membrane fluidity experiments

Membrane fluidity was assessed using the pyrenedecanoic acid probe (PDA), a probe analog to lipids that incorporates into the cell membrane. The probes in the membrane form monomers and excimers, with a rate of excimer proportional to the membrane fluidity. Under PUFAs (C18:2 n-6, C18:3 n-3, C22:6 n-3) application at 10 µM and 100 µM, we measured the emission spectrum of the PDA: 470 nm for the excimers and 400 nm for the the monomers. The monitoring of the fluorescent ratio 470/400 nm shift over time with PUFAs allowed a quantitative monitoring of the membrane fluidity changes due to PUFAs insertion into the membrane. Briefly, HEK/hTREK-1 cells were grown in culture in glass bottom culture dishes (MatTek, Ashland, U.S.A). Before fluorescent experiment, adherent HEK/hTREK-1 cells were incubated 1h at room temperature in the dark with 5 µM of the fluorescent lipid reagent probe in buffer provided in the membrane fluidity kit (Abcam). After incubation, the unicorporated probes were removed by washing cells twice with the same extracellular medium that used for WCR experiments. Epifluorescence microscopic experiments were performed on a 40X lens using excitation light filter with a 370-mm dichroic filter (Zeiss). Emitted light were taken at 480±15 nm and 405±10 nm and the 480/405 nm ratio was calculated at intial time before PUFA application (T_0_) and T_2_ (2 min), T_4_, T_6_, T_8_, T_10_, T_20_, T_30_, T_40_, T_50_ and T_60_. The 470/400 nm ratio was calculated as the average ratio overs 5 s. Then, the 470/400 nm ratio was normalized as (F/F_0_)-1 and represented over time (**Fig 7**) with F the ratio of fluorescence corresponding to a time T_t_ and F_0_ the ratio corresponding to the T_0_.

### Microsomes preparation

Microsomes were obtained from the HEK/hTREK-1 cell line and from the HEK 293T cell line as control. Briefly, cells were cultured in T75 flasks until confluence.Then, cells were washed with PBS twice and centrifugated 5 min at 15,000 RPM. The pellet was lysed in a 20 mM PIPES buffer (300 mM Sucrose, 20 mM PIPES, pH 7 with NaOH). Membranes were then mechanically braked up using insulin syringe. Samples were centrifuged 20 min at 10,000 g at 4°C. Lysis protocol was repeated twice. The pellet was discarded and the supernatant containing plasma membrane and thus transmembrane proteins was ultracentifugated 1h at 32,000 RPM at 4°C (Optima^TM^L-90K Ultracentrifuge, Beckamn Coulter; Rotor SW60Ti). The pellet containing microsomes was resuspended in 5 mM PIPES (300 mM Sucrose, 5 mM PIPES, pH 7.4 with NaOH) at a protein concentration adjusted to 25 mg/mL and conserved at −20°C 2 days before the spectral shift measurment.

### Spectral shift measurement

Microsomes of HEK 293T cell lin and HEK/hTREK-1 cell line were used to evaluate the TREK-1-PUFAs interaction with HEK 293T microsomes as control. Briefly, the proteins contained in the microsomes were labeled on the lysine residues with a fluorescent dye (MO_L011 RED-NHS 2^nd^ generation, NanoTemper Tehnologies GmbH) following the manufacturer protocol. The 20 µL at 600 µM of dye were used to labelled 200 µL of microsomes, thus leading to a final incubation volume of 200 + 20 µL. After 20 min of incubation, the 220 µL of labeled microsomes were applied to a gravity size exclusion B-column supplied in the kit, following the manufacturer protocol until etution. Labeled microsomes were eluted with 5 x 200 µL of the 5mM PIPES buffer. 5 fractions of 200 µL were collected in a clean Eppendorf. The first elution fraction contains the labeled microsomes while the last fractions contain more free dye. The fluorescence intensity of fraction 1, 2 and 3 were verified and fractions with a fluorescence count between 400 and 2,000, using 100% excitation power, were used for the experiment after a 10 min centrifugation at 16,000 g to avoid homogeneity problem. A serial dilution of C22:6 n-3 over 16 points following 1:1 pattern was set up and subsequently mixed with labeled microsomes and loaded into standard capillaries (NanoTemper Technologies GmbH). The total proteins concentration in microsomes was kept around 2,5 µg/mL whereas C22:6 n-3 was titrated from 1 mM to 30,5 nM. The read out was performed on a Monolith X instrument from NanoTemper Technologies GmbH. The samples were subjected to SpS and MST measurments and only SpS was analyzed due to the normalization requirments. The changes in the maxima emission wavelength following a ratiometric measurement upon protein-C22:6 n-3 complex formation was used to generate a binding curve as a function of C22:6 n-3 concentrations (Langer *et al*, 2022). The data were pulled from 9 and 5 individual repeat for the TREK-enriched microsomes and from HEK 293T microsomes respectively. Data processing and analysis were carried out using the MO.Control 3 software from NanoTemper Technologies followed by normalization as described in the results section. All the data points presenting signs of irregularity were automatically discarded upon merging the data, as documented in the software.

### Chemicals

In this study we tested one direct activator of TREK-1, ML402, one saturated fatty acid, stearic acid, one monounsaturated fatty acid, oleic acid, 9 different PUFAs having between 18 and 22 carbons and 2 to 6 double bonds (**Table 1**). All reagents are summarized in the reagents and tools table. 10 mM stock solutions were prepared by dissolving PUFAs in absolute ethanol, ML402 in DMSO. PUFAs and ML402 were stored at −80 °. 10 µM solutions were obtained by diluting an aliquot of stock solution in the extracellular medium just before use for WCR experiments. Cells for current recordings were first superfused with extracellular medium containing ethanol at 1/1000 as a control.

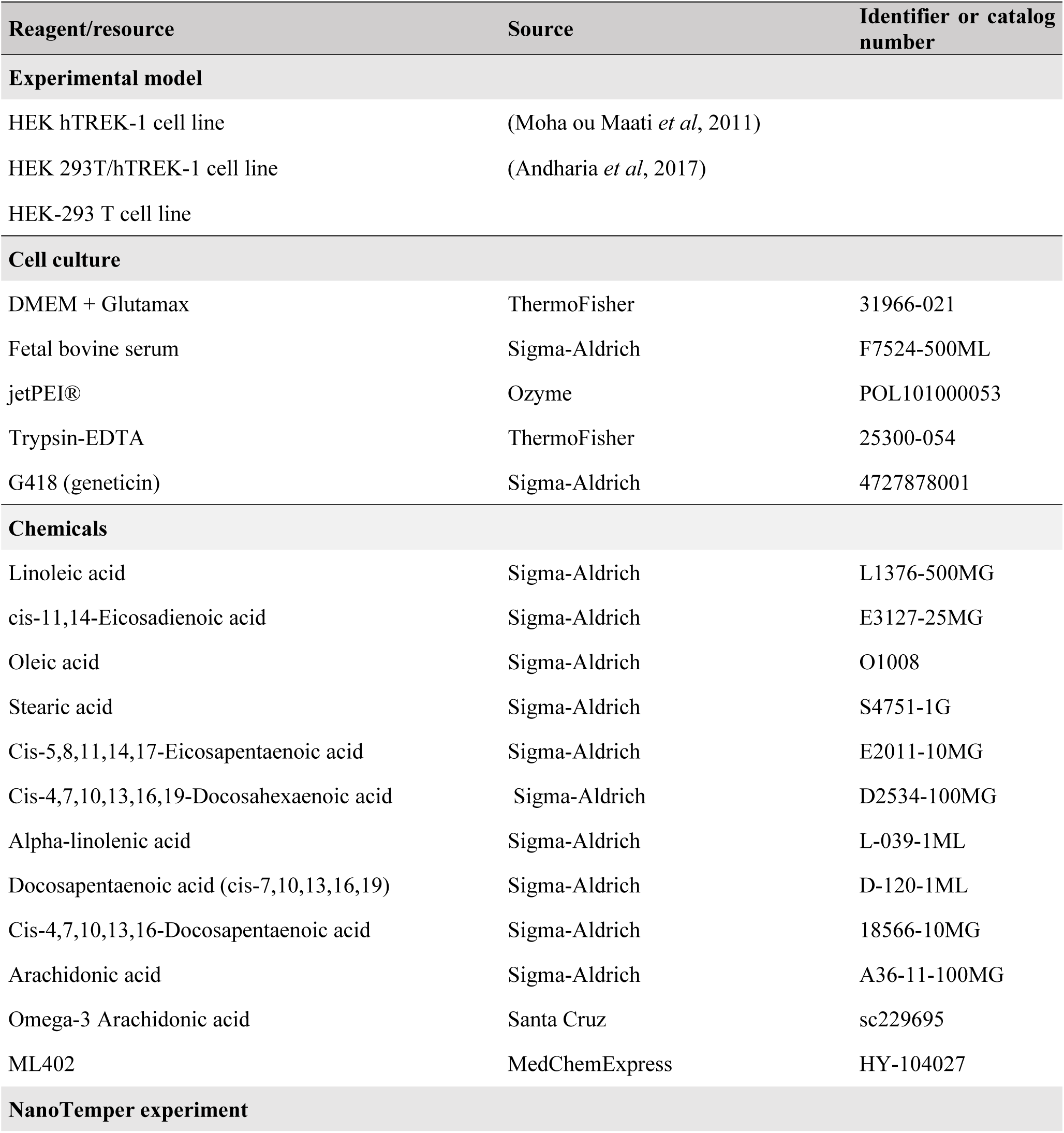
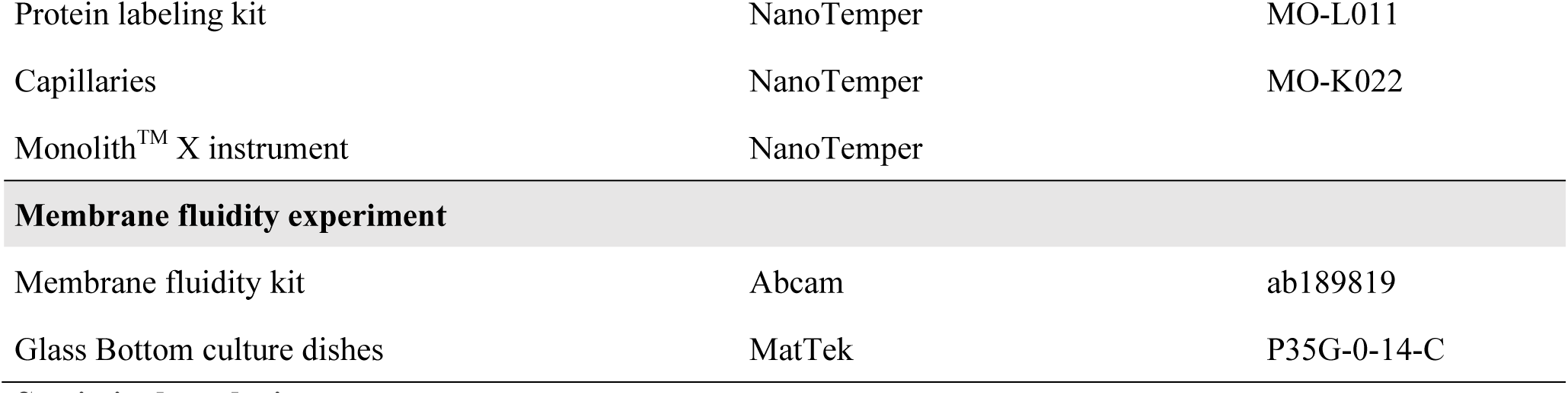

### Statistical analysis

All set of experiments were performed on at least three different batches (congelation and/or passage) of cells. All descriptive statistics are displayed from **Table 1** to **Table 4**. Statistical analysis were performed using Prism software (GraphPad Prism 9, Inc., USA). Spearman correlation test were performed and the p-value indicates the significance of the correlation and the r parameter indicates the direction of the correlation (negative or positive). For the linear regression, the p-value indicates the significance of the relationship, the R^2^ indicates the goodness of the fit an the error bars represent the 90 % confidence interval. Kruskal-Wallis test followed by a *post hoc* Dunn’s test was used for multiple comparison of unmatched data. A p-value of 0.05 or less was considered as statistically significant. The differences between more than 2 groups are displayed by the letter code, where groups that do not share the same letter are significantly different.

## AUTHORS CONTRIBUTION

EB carried out the experiments (WCR and IO), set up experiments and protocols, conceived the original idea and wrote the manuscript.

EA carried out the experiments (WCR) and discussed the results.

JB carried out the experiments (WCR) and discussed the results. JL carried out the experiments(WCR) and discussed the results. XB carried out the experiments (IO) and discussed the results.

AE participated to the PUFAs structure discussion.

CV helped supervise the project on chemistry of PUFAs and discussed the results.

PS helped in the design and result analyses of affinity tests.

TD helped supervise the project on chemistry of PUFAs and discussed the results.

JMG helped supervise the project on chemistry of PUFAs and discussed the results.

CO helped supervise the project on chemistry of PUFAs and discussed the results.

JYL set up experiments and protocols, conceived the original idea, wrote the manuscript and supervised the project.

HMM provided HEK hTREK-1 cells

MD set up experiments and protocols, conceived the original idea, wrote the manuscript and supervised the project.

## ACKNOWLEDGEMENT

This work is supported by the Agence Nationale de la Recherche: acronym TA-BOOm (ANR-20-CE44-0001), Région Occitanie Pyrénées-Méditerranée (MNE-AGPI03), Key Initiative Montpellier Université d’Excellence (KIM Biomarkers and Therapies) and the CBS2 doctoral school. We thank Pr OKADA for the kind gift TR-1 cells: a 293 T cell line that stably expresses TREK-1 channel.

## Notes

### Competing Interest Statement

The authors have declared no competing interest.

### Summary of Updates

All the paper is update, but the title

